# A simple active fluid model unites cytokinesis, cell crawling, and axonal outgrowth

**DOI:** 10.1101/2024.05.22.595337

**Authors:** Erin M. Craig, Francesca Oprea, Sajid Alam, Ania Grodsky, Kyle E. Miller

## Abstract

Axonal outgrowth, cell crawling, and cytokinesis utilize actomyosin, microtubule-based motors, cytoskeletal dynamics, and substrate adhesions to produce traction forces and bulk cellular motion. While it has long been appreciated that growth cones resemble crawling cells and that the mechanisms that drive cytokinesis help power cell crawling, they are typically viewed as unique processes. To better understand the relationship between these modes of motility, here, we developed a unified active fluid model of cytokinesis, amoeboid migration, mesenchymal migration, neuronal migration, and axonal outgrowth in terms of cytoskeletal flow, adhesions, viscosity, and force generation. Using numerical modeling, we fit subcellular velocity profiles of the motions of cytoskeletal structures and docked organelles from previously published studies to infer underlying patterns of force generation and adhesion. Our results indicate that, during cytokinesis, there is a primary converge zone at the cleavage furrow that drives flow towards it; adhesions are symmetric across the cell, and as a result, cells are stationary. In mesenchymal, amoeboid, and neuronal migration, the site of the converge zone shifts, and differences in adhesion between the front and back of the cell drive crawling. During neuronal migration and axonal outgrowth, the primary convergence zone lies within the growth cone, which drives actin retrograde flow in the P-domain and bulk anterograde flow of the axonal shaft. They differ in that during neuronal migration, the cell body is weakly attached to the substrate and thus moves forward at the same velocity as the axon. In contrast, during axonal outgrowth, the cell body strongly adheres to the substrate and remains stationary, resulting in a decrease in flow velocity away from the growth cone. The simplicity with which cytokinesis, cell crawling, and axonal outgrowth can be modeled by varying coefficients in a simple model suggests a deep connection between them.

## INTRODUCTION

Axonal outgrowth occurs when the growth cone moves forward. It is supported by the transport of membrane-bound organelles, slow axonal transport of soluble proteins, and bulk flow of polymerized cytoskeletal elements (Miller and Suter, 2018; Pfenninger, 2009; Roy, 2020). Collectively, these mechanisms move proteins, mainly made in the cell body, into and along the axon. Similarities between morphology, patterns of cortical flow, and gene conservation have motivated many groups to make mechanistic analogies between cytokinesis, cell crawling, and axonal outgrowth (Bray and White, 1988; Fritz-Laylin, 2020; Karsenti and Nedelec, 2004; Michaud et al., 2021; Miyata et al., 2020; Mogilner and Craig, 2010). Nonetheless, the relationships between these modes of motility are poorly understood. To better understand axon outgrowth, we explore its biophysical similarities and differences with cytokinesis and cell crawling.

Bulk cytoskeletal flow underlies cellular motion. It involves the movement of crosslinked arrays of microtubules, actin filaments, intermediate/neurofilaments, and embedded organelles (Miller and Suter, 2018). It is a eukaryotic innovation driven by molecular motors and cell adhesion molecules, including non-muscle myosin II (NMII) and integrins (Richards and Cavalier-Smith, 2005; Sebe-Pedros et al., 2010). It underlies the processes of cytokinesis (Singh et al., 2019), mesenchymal migration (Schaub et al., 2007), amoeboid migration (Bergert et al., 2015), neuronal migration (He et al., 2010), and axon outgrowth (Athamneh et al., 2017). The mechanisms driving bulk flow are best understood during mesenchymal migration, where adherent cells crawl over two-dimensional surfaces (**Fig. 1A**). In this process, a convergence/transition zone enriched in non-muscle myosin II (NMII), along with actin assembly at the leading edge, drives retrograde actin flow. As a result of substrate interactions, traction forces are generated that pull the substrate rearward and the microtubule-rich cell body forward (Munevar et al., 2001; Salmon et al., 2002; Svitkina et al., 1997). Across a migrating cell, this creates a velocity profile such that the cell body moves forward at the rate of cell migration, and the lamellipodium is either stationary or moves rearwards depending on adhesion strength. Coupled with this, the actin meshwork is disassembled across the convergence zone (Vallotton et al., 2004). This generates soluble proteins, which are transported to the leading edge to support actin and adhesion assembly (Zicha et al., 2003). Through bulk flow, cytoskeletal assembly, and disassembly, the cell body, the position of the convergence zone, and the cell’s leading edge move forward on average in tandem in migrating cells.

**Figure 1.**
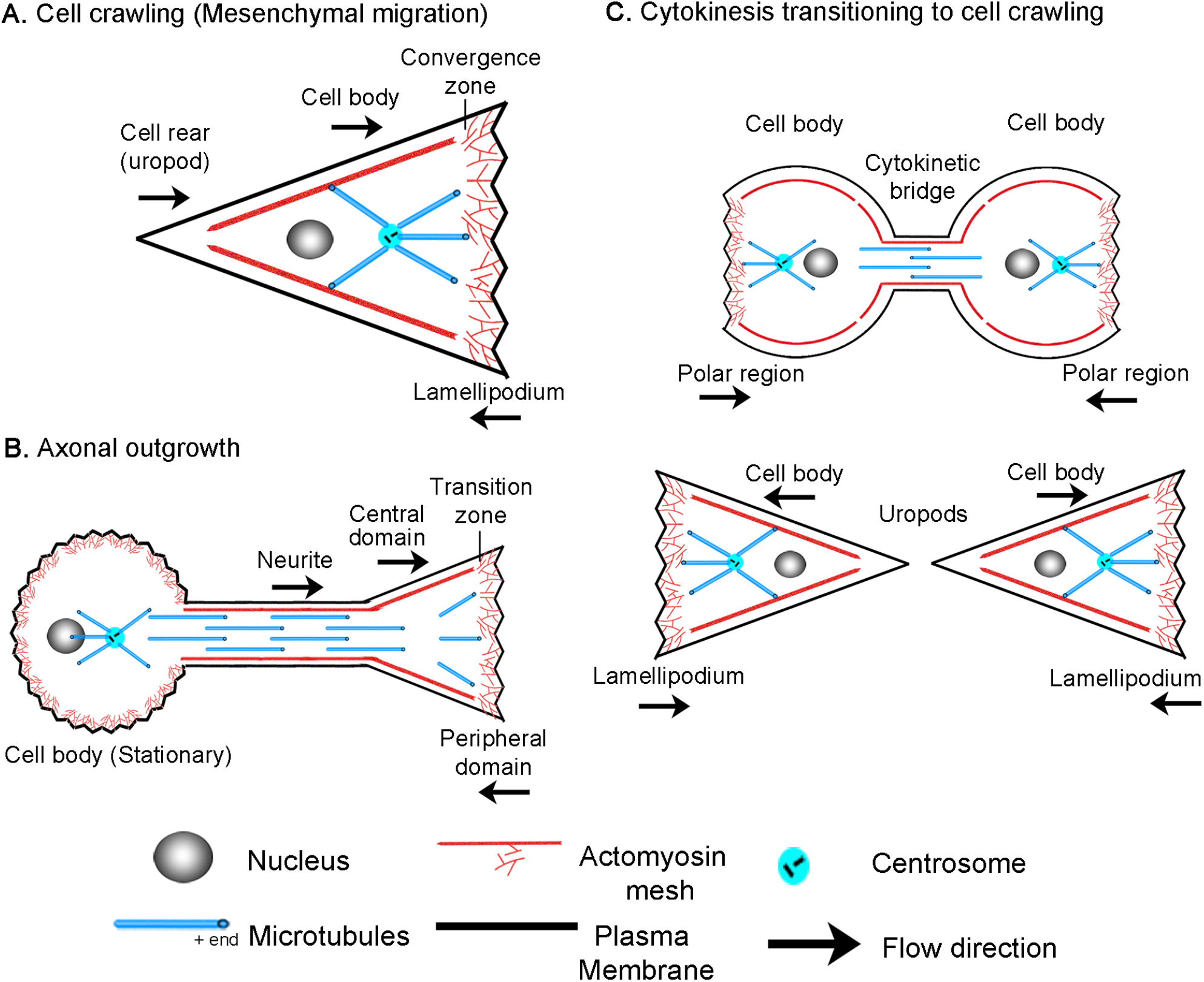
Overview of (A) cell crawling, (B) axonal outgrowth, and (C) the transition from cytokinesis to cell crawling following abscission.

Similarly, growth cones exhibit flow patterns akin to those of migrating cells (**Fig. 1B**), though comparable regions have a different nomenclature (Miller and Suter, 2018). In neurons, the lamellipodium is often called the peripheral domain (P-domain) and, like in crawling cells, exhibits retrograde actin flow and generates traction forces (Betz et al., 2011; Chan and Odde, 2008; Koch et al., 2012). This flow is driven by a combination of actin assembly at the leading edge and NMII-driven contraction at the convergence or transition zone (T-zone) (Medeiros et al., 2006; Rochlin et al., 1995). Directly behind is the organelle and microtubule-rich growth cone central domain (C-domain), which corresponds to the non-neuronal cell body. Like in crawling cells where MTs in the cell body move forward as the cell advances (Salmon et al., 2002), cytoskeletal elements and organelles in axons flow forward in bulk during outgrowth (Athamneh et al., 2017; Miller and Sheetz, 2006; Reinsch et al., 1991) and neuronal migration (He et al., 2010). Axonal outgrowth differs from neuronal migration and cell crawling in that along the axon, adhesions to the substrate dissipate forces (O’Toole et al., 2008). As a result, anterograde bulk flow decreases with distance from the growth cone, and the neuronal cell body is stationary.

Additionally, it has long been appreciated that cell crawling appears to be a continuation of cytokinesis (Bray and White, 1988; DeBiasio et al., 1996; Swann and Mitchison, 1958). In dividing cells, NMII activity at the cleavage furrow drives constriction to help form the cytokinetic bridge. Paired with substrate adhesions under the polar regions, traction forces are generated that pull daughter cells apart during the final steps of cytokinesis (Dix et al., 2018; Taneja et al., 2019) (**Fig. 1C**). After cells complete abscission, the cytokinetic bridge becomes the rear (uropod) and the polar regions transition into the lamellipodia of the daughter cells. Likewise, many genes essential for cytokinesis mediate cell crawling and axon outgrowth. For example, the Rap1/Rac/Cdc42 signaling axis is essential for controlling cell adhesion during cytokinesis, cell crawling, and axon outgrowth (Chircop, 2014; Dao et al., 2009; Govek et al., 2005; Ridley, 2015; Shah and Puschel, 2016). Whereas, the RhoA pathway regulates NMII activity, which is the primary forcing-generating motor in all three modes of motility. Furthermore, it is well appreciated that genes important for cell division ‘moonlight’ in axon outgrowth (Baas, 1999; Lu and Gelfand, 2017). For example, RhoA and Aurora A kinase, which regulate actin and MT dynamics, are activated at the site of axonal initiation *in vivo* (Pollarolo et al., 2011). Septins, which help specify the site of the cleavage furrow, are concentrated axonal branch points at the base of dendritic spines (Falk et al., 2019), and the mitotic motors, Kinesin-5 (Kahn et al., 2015), Kinesin-12 (Liu et al., 2010), and Kinesin-14 (Muralidharan and Baas, 2019) modulate the rate of axon outgrowth and axonal initiation. While genes and mechanisms that power cytokinesis, cell crawling, and axon outgrowth are often shared, the mechanistic links between these processes are poorly understood.

To address this, here, we develop an active fluid model (de Rooij et al., 2017; Julicher et al., 2007; Marchetti et al., 2013; Mogilner and Manhart, 2018) to describe flow patterns and motion during cytokinesis, mesenchymal migration, amoeboid migration, neuronal migration, and axon outgrowth. Noting that these processes can be described by a single model by varying the size and position of the convergence zone and the profile of subcellular adhesions suggests the biophysical mechanisms underlying axonal outgrowth may be directly related to cell crawling and cytokinesis.

## RESULTS

To develop a biophysical intuition for the commonalities and differences between bulk flow during different modes of cell motility, we present a side-by-side comparison using data previously collected from our own and other labs (Athamneh et al., 2017; Bergert et al., 2015; He et al., 2010; Schaub et al., 2007; Singh et al., 2019) (**Fig. 2**). Before reviewing the data, it is essential to point out that while cells are often drawn as bags of fluid containing a few isolated cytoskeletal elements, stunning electron microscopy images reveal they are densely packed with a cross-linked meshwork of cytoskeletal elements and embedded organelles (Hirokawa, 1982; Vassilopoulos et al., 2019). In particular, supporting the plasma membrane is a cortical meshwork of actin, NMII, alpha-actinin, and spectrin, which is directly linked to crosslinked MTs through proteins such as spectraplakins (Svitkina, 2018; Verkhovsky et al., 1995; Voelzmann et al., 2017). Based on studies in migrating fibroblasts and growth cones, soon after MTs assemble, they form crosslinks with the actin meshwork (Salmon et al., 2002; Schaefer et al., 2002). As a result, MTs and actin filaments have similar velocities and flow patterns. Likewise, in neurons, beads bound to the outside of axons, docked mitochondria, microtubules, and phase-dense objects all move with similar velocity profiles during elongation (Athamneh et al., 2017; Lamoureux et al., 2010a). Because the cytoskeletal meshwork and embedded organelles are tightly cross-linked, in general, tracking the motion of a given component such as speckle-labeled actin filaments, NMII, alpha-actinin, speckle-labeled microtubules or docked mitochondria provides information about the overall bulk flow of the meshwork.

**Figure 2.**
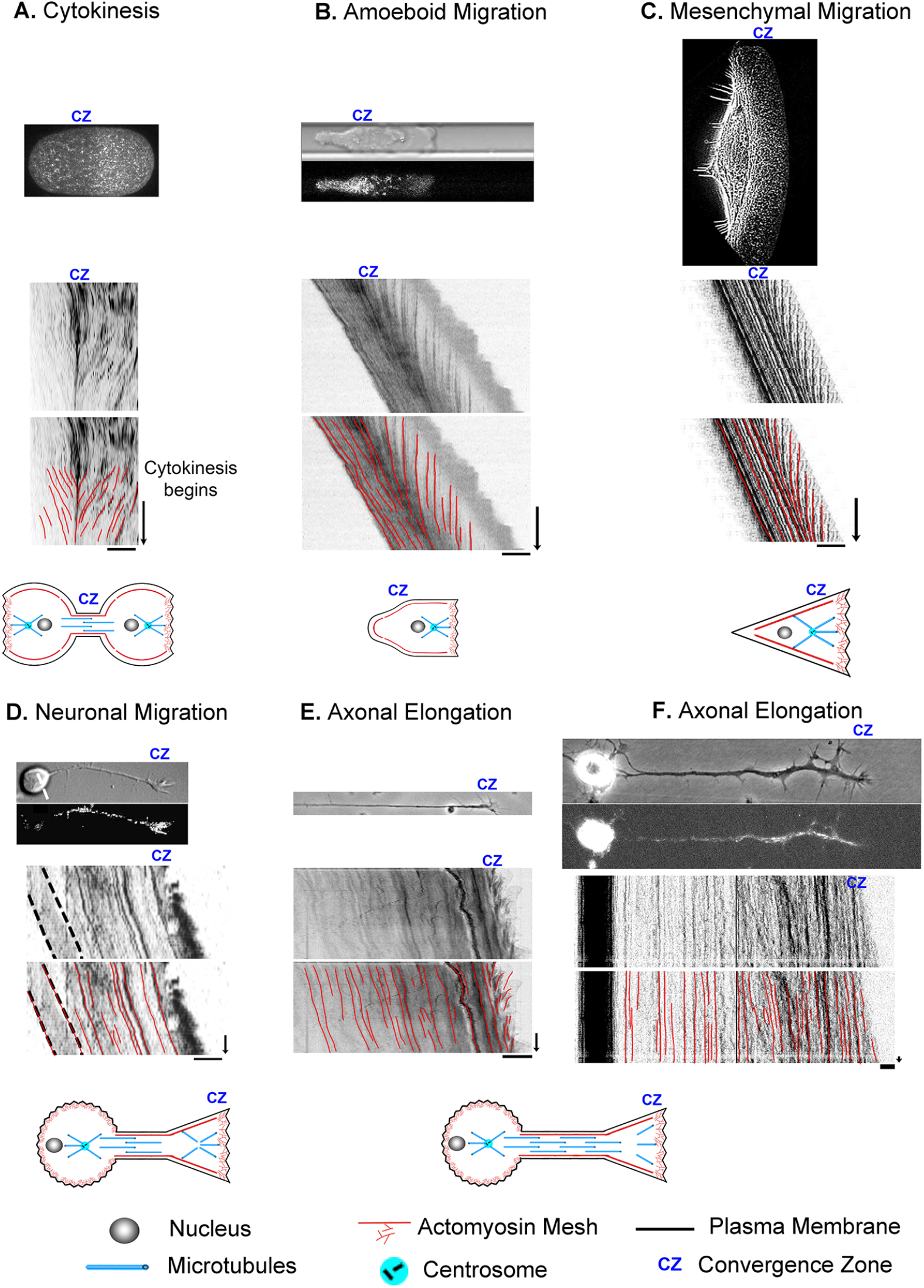
Cell division, motility, and axon outgrowth share common features of motion. (A) Cytokinesis in a *C. elegans* embryo. The top image in each panel shows a still image of the cell, with unmarked and marked kymographs and schematics below to illustrate the cell type and position of the convergence zone. (B) Amoeboid cell migration of a constrained Walker carcinoma cell. (C) Mesenchymal migration of a fish keratocyte. (D) Neuronal cell migration of a rat cerebellar granule cell. (E) Axonal outgrowth of chick sensory neurons imaged with phase microscopy to track retrograde flow in the growth cone and (F) by fluorescent microscopy of mitochondria to track bulk axon motion along proximal axon and cell body. In all panels, the bar = 10 *μ*m and the arrow = 1 min.

With this background, cytokinesis and amoeboid migration share a substantial similarity in morphology and subcellular flow patterns. In both, there is bilobate morphology with a continuous flow toward a convergence zone (Basant and Glotzer, 2018; O’Neill et al., 2018). To illustrate, we show a kymograph of cytokinesis during the first cell division in an unconstrained wild-type *C. elegans* embryo expressing non-muscle myosin II GFP (Singh et al., 2019) (**Fig. 2A)**. At the onset of cytokinesis, which occurs halfway down the time axis of the kymograph, actomyosin flows symmetrically towards a convergence zone at the cleavage furrow at roughly 6 *μ*m/min while the cell remains stationary. Similarly, during amoeboid migration of constrained Walker carcinoma cells expressing non-muscle myosin II regulatory light chain GFP, there is an inward flow toward a convergence zone in the middle of the cell. It differs from cytokinesis in that across the leading edge, material flows slowly rearward at a rate of (−0.4 *μ*m/min), and instead of being stationary, the cell body flows rapidly forward at a rate of 7.9 *μ*m/min (**Fig. 2B**) (Bergert et al., 2015; Liu et al., 2015). In a similar manner, during mesenchymal migration of fish keratocytes labeled with rhodamine-phalloidin to track actin filaments (Schaub et al., 2007), rapid advance of the cell body (12 *μ*m/min) is paired with slow retrograde flow (−1 *μ*m/min) at the leading edge. (**Fig. 2C**). While neurons have dramatically different morphologies, analysis of neuronal migration of rat cerebellar granule cells labeled with alpha-actinin-GFP to track actin filaments indicates that, like non-neuronal cells, the cell body and leading process advance in unison (3 *μ*m/min) and retrograde flow occurs in the growth cone P-domain (−1.5 *μ*m/min) (He et al., 2010) (**Fig. 2D**). The major difference is that the convergence zone is much shorter in length. The flow pattern in the distal axons of sensory neurons, a type of neuron found in the peripheral nervous system, appears to be similar to flow patterns in neuronal migration (**Fig. 2E, F**). Like neuronal migration, the convergence zone is positioned in the growth cone, and across the P-domain, retrograde flow occurs at -3 *μ*m/min. Where neuronal migration and outgrowth differ is that while materials flow forward in bulk in the distal axon at a peak rate of 0.5 *μ*m/min, because the cell body is stationary, flow velocity decreases with distance from the growth cone (Athamneh et al., 2017; Miller and Sheetz, 2006). Collectively, the flow patterns suggest these processes are related but differ in subcellular patterns of force generation, viscosity, and substrate adhesion. Motivated by this observation, our goal was to determine if these diverse forms of motility can be united in a single simple mathematical model, where variations in the profiles of sub-cellular adhesion and internal force generation explain the differences in motility.

### Developing a generalized model of cell motility

To model bulk flow, we use an “active Maxwell fluid” approach, which in its complete form considers both solid-like and fluidlike behaviors, internal force generation, and substrate adhesions (Craig et al., 2015; de Rooij et al., 2017; Marchetti et al., 2013; Rubinstein et al., 2009). At a molecular level, this theory treats cells as collections of filaments connected by dynamic cross-linkers. Over short periods, a cell behaves as a solid because cytoskeletal elements are cross-linked. Yet, over longer periods, cells act as fluids that flow in response to forces. The reason materials act as solids over short times and fluids over long times arises through the dynamics of the molecular interactions within a material. A concrete way to understand this is to consider two filaments held together with multiple dynamic spring-like crosslinkers (de Rooij et al., 2017). If both filaments are stationary, a subset of crosslinkers will bind both filaments based on the association and disassociation rates of the binding interactions. When force is applied abruptly, the molecular interactions between the components don’t have time to break. Thus, the material behaves as a purely elastic material with parameters determined by the number and the spring constants of the bound crosslinkers. Yet, if a continuous force is applied, the filaments slide apart at a constant average rate determined by the spring constants and dynamics of the crosslinkers. This occurs because, at a microscopic level, when a single crosslinker unbinds, the force is transferred to the remaining crosslinkers, and filaments slide apart due to the stretching of the attached springs. When a crosslinker remakes a connection, it is initially unstressed, but as the filaments slide apart, it also stretches. Whether a time period is considered short or long depends on the kinetics of the crosslinkers. At a macroscopic level, this is defined by the term, *τ*, given in units of time, which mathematically equals the ratio of viscosity (*μ*) over elasticity (*E*) (i.e., *μ* = *E τ*). Experimentally, the *τ* for neurons is on the order of 10 seconds in the growth cone (Betz et al., 2011) and 8.5 minutes in *Drosophila* nerves (Purohit, 2015). Because we model cellular behaviors occurring continuously, cells behave as fluids on relevant time scales, and the elastic terms can be dropped.

What makes cells an “active” rather than passive fluid are the molecular motors and cytoskeletal dynamics powered by ATP consumption (Julicher et al., 2007). When force generation occurs at a higher level locally, for example, at a convergence zone, it drives the surrounding flow of material. Following the approach of other active fluid models for cellular mechanics (Craig et al., 2015; de Rooij et al., 2017), we assume the cytoskeleton maintains a steady-state structure with cellular elastic properties negligible over our time scale of interest and write the balance of motor-driven active stress, viscous stress, and traction that governs the flow of cytoskeletal material as:

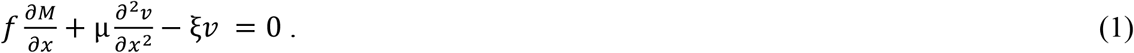

The first term in eq. 1 corresponds to contractile stress from motor proteins, which is proportional to the gradient of motor density, 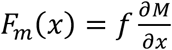. This equation states that while motors may be active throughout the cell, net force vectors only arise over regions where motor activity varies over distance. The second term corresponds to internal viscous stress arising through cell deformation, 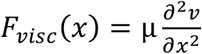, where *μ* is the viscosity and *ν*(*x*) is the local cytoskeletal flow rate. It states that internal deformation, represented by 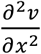, is highest for large viscous stress, *F*_*visc*_(*x*), and low viscosity, *μ*. The third term represents external traction, *T*(*x*) = *ξν*, which we write as being proportional to the velocity of local cytoskeletal flow (*ν*) and the strength of adhesions (*ξ*) (Chan and Odde, 2008). In cases where cells are not attached to the substrate, such as in *Dictyostelium* cultured in suspension, the traction force is equal to zero, and flow occurs at a maximal rate determined by the interaction of force generation and the viscosity terms in Eq. 1. In turn, when internal force generation (*f*) and viscosity (*μ*) are held constant, as adhesion strength increases the velocity of flow decreases and traction forces increase as described by the substrate-coupling / clutch hypothesis (Mitchison and Kirschner, 1988; Suter et al., 1998). Because the experimental data suggest the effective adhesion coefficients on the right and left sides of the convergence zone are asymmetrical, we define them as *ξ*_*R*_ and *ξ*_*L*_, respectively, where we adopt a coordinate system in which the leading edge of the cell is to the right of the convergence zone. (See Table 1 for a summary of the model parameters and terminology.) For generality, we assume that force generation inclusively arises from diverse mechanisms, and substrate adhesions arise through specific and non-specific interactions (Bell et al., 1984; Clarke and Martin, 2021). Careful examination of the nano-scale architecture of adhesions indicates that proteins such as alpha-actinin, which are directly coupled with actin, move at the highest rates during retrograde flow and that proteins binding to the extracellular matrix, such as integrins, move significantly slower (Case and Waterman, 2015). While this could be modeled by considering the motion and physical parameters of each distinctive molecular layer (e.g., actin, alpha-actinin, vinculin, talin, integrins, and laminin), here we operationally define adhesions as a single layer linking the actin cytoskeleton and the extracellular matrix.

**Table 1:** Definitions of model parameters and variables.

| Model variables and parameters | Units | Significance |
| --- | --- | --- |
| $E$ | $Pa$ | Young's Modulus |
| $f$ | $pN$ | Force per motor |
| $F_m(x)$ | $pA$ | Local stress induced by motor proteins |
| $F_{max}$ | $pA$ | Maximal stress at position $x_c$ |
| $F_{Normal}$ | $N$ | Normal force |
| $F_{visc}(x)$ | $pA$ | Local viscous stress |
| $g$ | $m/s^2$ | Gravitational acceleration |
| $L$ | $\mu m$ | Length of cell (from cell rear to leading edge) |
| $\lambda \equiv \frac{2\mu}{\sigma^2 \xi}$ | *dimensionless* | Dissipation length relative to motor width distribution |
| $\sqrt{\frac{\mu}{\xi}}$ | $\mu m$ | Characteristic dissipation length |
| $M_0$ | $\mu m^{-1}$ | Maximum motor density |
| $M(x)$ | $\mu m^{-1}$ | Linear density of motor proteins |
| $\sigma$ | $\mu m$ | Width of motor distribution |
| $\tau$ | $s$ | Time constant |
| $T(x)$ | $pA$ | Traction stress |
| $T_{Rope}$ | $N$ | Tension on rope |
| $TF_{Gravel, Ice}$ | $N$ | Traction forces |
| $\mu$ | $\frac{pNs}{\mu m}$ | Viscosity |
| $\mu_{ice, gravel}$ | *dimensionless* | Friction coefficients |
| $v$ | $\mu m/s$ | Velocity |
| $x$ | $\mu m$ | Distance along cell axis |
| $x_c$ | $\mu m$ | Maximal $F_m(x)$ location |
| $x_{CZ}$ | $\mu m$ | Position of convergence zone |
| $\xi_{L,R}$ | $\frac{pAs}{\mu m}$ | Adhesion strength, left or right of convergence zone |

To develop a general model of cell migration, we simplify the cell into three regions: A convergence zone in the middle that generates contractile forces, a leading edge (shown to the right in **Fig. 3B**), and a lagging region (shown to the left in **Fig. 3B**). Depending on the context, the leading edge can represent actomyosin flow on the right side of a cell undergoing cytokinesis, the front of cells during amoeboid migration, or the lamellipodial/filopodial region of cells undergoing mesenchymal migration, neuronal migration, or axonal outgrowth. In all cases, the leading edge has a zone of actin filament assembly at the leading-edge boundary and disassembly at the convergence zone. Since we consider steady-state conditions, the length and mass of the area are constant over time as the result of coupled transport of soluble cytoskeletal elements from sites of disassembly to assembly (Zicha et al., 2003). Likewise, the lagging edge can represent the left side of a cell undergoing cytokinesis, the rear of the cell body and trailing uropod of amoeboid cells, the cell body and surrounding cytoskeletal elements of migrating mesenchymal and neuronal cells, or the distal axon and cell body of neurons. Except for cytokinesis, where the left and right sides are symmetric, the lagging edge represents the interconnected meshwork of actomyosin, microtubules, associated cross-linkers, motors, and organelles. While contraction and extension can occur over the lagging edge, for migrating cells under steady-state conditions, the length remains constant, and flow in the lagging edge represents cell motion. The one exception is that during axon outgrowth, we define the left boundary of the lagging edge as the stationary cell body and the advance of the right boundary as reflecting axonal lengthening.

**Fig 3.**
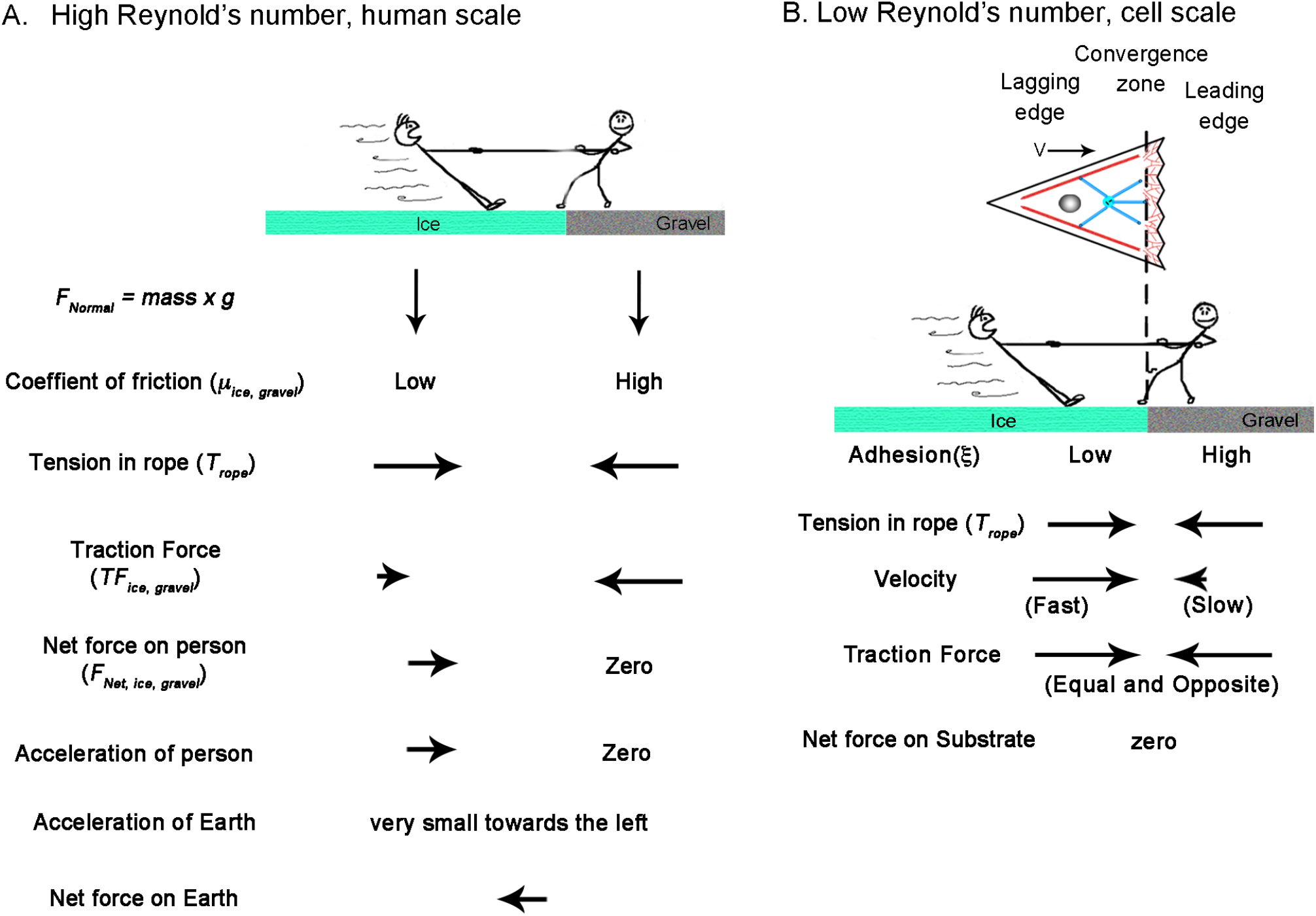
How Reynolds number affects the physics of a Tug-of-War. A. Tug of war at the human scale where the Reynolds number is high. The balance of forces accelerates the person on ice to the right, which is balanced by the acceleration of the earth to the left. B. A cellular tug of war at a low Reynolds number illustrating mesenchymal migration. Here, acceleration terms become negligible, and traction force is a function of velocity. The dotted line indicates the position of the convergence zone, with strong adhesions to the right and weaker adhesions to the left. While the center of mass shifts to the right, the net traction forces on the leading and lagging sides are balanced; thus, the net force on the substrate is zero.

As to how this model gives rise to motion, it notably diverges from classic approaches (DiMilla et al., 1991) in that it assumes the traction forces on the leading and lagging sides are always equal and opposite and that gradients in internal flow fields that shift the center of mass, rather than traction force imbalance, drives migration. This may seem problematic for creating motion, as there is no net force vector acting on the substrate, but it is explained by considering how inertial forces and friction differ under high and low Reynolds number conditions (**Fig. 3**). Doing so also illustrates how human intuition sometimes leads to misconceptions when applied to cellular biophysics.

From the viewpoint of a stationary reference frame, consider a high Reynold’s number macroscopic tug of war between two people standing on the Earth (**Fig. 3A**). They have the same mass and normal force pushing them downwards (*F*_*Normal*_ = *g* × *mass*). One stands on ice and the other on gravel, with correspondingly low and high friction coefficients (*μ*_*ice*_, *μ*_*gravel*_). Through muscle contraction, a moderate level of tension acts on the rope (*T*_rope_), so the person on the ice slides, but the person on the gravel remains stationary. When the traction force under the person on the gravel (*TF*_*gravel*_) is measured, it is equal to the tension on the line. In contrast, the traction force under the person on ice (*TF*_*ice*_) equals their normal force multiplied by *μ*_*ice*_ according to Amontons’s law. This difference between tension and traction force results in the net force. Keeping in mind that the two people represent the front and back of a cell, dividing their net mass by net force yields their acceleration. Accordingly, to maintain force balance, the earth accelerates in the opposite direction by a small amount. Thus, if cells behaved like macroscopic objects, their motion would generate a net force on the substrate.

Moving to a microscopic low Reynolds number scale, two differences arise (**Fig. 3B**). The first is that because of viscous drag, constant force results in a constant velocity and, thus, zero acceleration. Secondly, instead of the traction force being a constant determined by the friction coefficient and the normal force, it depends on the product of the adhesion coefficient and velocity. Consequently, because tension on the rope pulls the two people with an equal force, the person on the ice slides rapidly to the right, while the person on the gravel slides slowly to the left. Because this results in equal and opposite traction forces, the net force on the substrate is zero. Collectively, motion occurs because the net movement of ‘people’ to the right is greater than to the left.

To express this idea more formally, we project three-dimensional cells to a 1D domain extending from the rear (*x* = 0) to the leading edge (*x* = *L*), with the site of maximum contractile force generation denoted as *x* = *x*_*CZ*_ (**Fig. 4A**). We write the steady-state motor distribution, *M*(*x*), as a Gaussian function of width *σ* and maximum motor density *M*_0_, centered on the location of the convergence zone *x*_*CZ*_:

**Figure 4.**
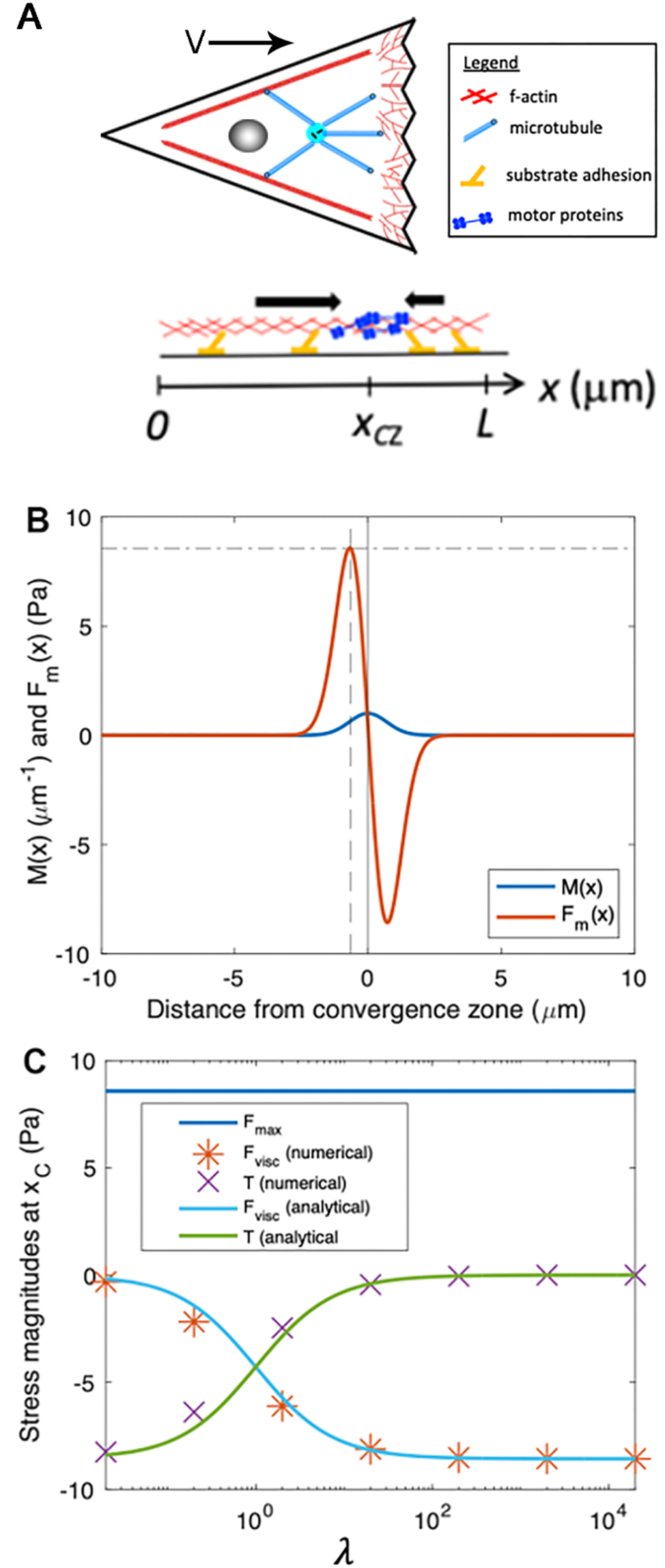
Numerical modeling generates output that matches analytic predictions. (**A**) Schematic of biophysical model for force generation and adhesion distributions within a cell, projected onto one dimension, where we define a coordinate system with *x* = 0 as the rear of the cell, *x* = *L* as the leading edge of the cell in the direction of migration, and *x* = *x*_*CZ*_ as a convergence zone where motor-based contractility is localized. Note that while the cartoon only shows motors which are enriched at the CZ, they are distributed throughout the cell. **(B)** Motor protein density (Eq. 2) and corresponding motor-induced stress (Eq. 3) for parameters: *L* = 20*μm, x*_*CZ*_ = 10*μm, σ* =1*μm*, 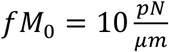. The vertical solid line represents the location of the convergence zone, *x*_*CZ*_. The vertical dashed line corresponds to 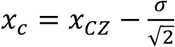, where motor-induced stress is maximized, and the horizontal dashed line corresponds the maximum motor-induced stress, *F*_*max*_. **(C)** *F*_*max*_, *F*_*visc*_, and *T* at location *x*_*c*_ as a function of *λ*. Analytical approximations for *F*_*visc*_ (Eq. 7) and *T* (Eq. 8) are shown with solid lines. The data markers (^*^ and ×) correspond to numerical solutions to the stress balance equation (Eq. 1).

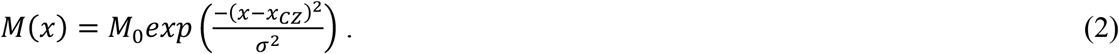

From this, it follows that the contractile stress from motor proteins, *F*_*m*_(*x*), is proportional to the gradient of motor density

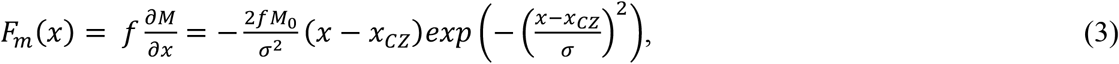

where *f* is the contractile force per motor. Using the steady-state assumption that forces from leading-edge actin polymerization and the corresponding membrane tension are closely balanced (Miller and Suter, 2018; Sens and Plastino, 2015), we adopt the boundary conditions that velocity is uniform at the cell edges: 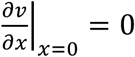 and 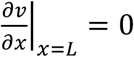. Simply stated, this indicates gradients in cytoskeletal flow approach zero near the boundaries.

To illustrate the basic features of our minimal cell model, **Fig. 4B** shows example plots of *M*(*x*) and *F*_*m*_(*x*) (Eqs. 2 and 3). Because the stress is proportional to the derivative of motor distribution, it exhibits two peaks such that cytoskeletal material is pulled inward toward the convergence zone. The location of these local maxima of *F*_*m*_(*x*), denoted as *x*_*c*_, can be determined by the condition: 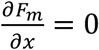, yielding: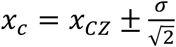. The magnitude of the maximum motor-induced stresses is given by: 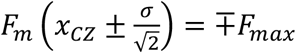, which simplifies to 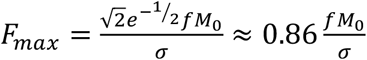. Collectively this model relates the distribution of the motors, viscosity, and adhesion with internal patterns of force generation, flow, and migration speed.

### Estimating stress and flow speed near the convergence zone

By numerically solving Eq. 1 for the velocity profile, *ν*(*x*), we can predict the distribution of stresses and flows across the length of the cell for a set of physical parameters. First, to develop intuition, we make analytical estimates of the flow speeds and relative magnitudes of *F*_*m*_(*x*), *F*_*visc*_(*x*), and *T*(*x*) at the location of maximum motor-based stress *x*_*c*_ (**Fig. 4B**).

To estimate the passive internal viscous deformation at a location, we use the numerical approximation:

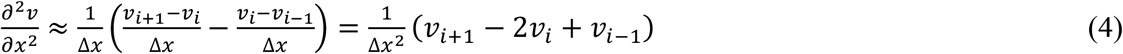

where we consider values of internal flow speed at discrete locations (*i*.*e*., the indexes i, i+1, and i-1) separated by a small spatial increment, Δ*x*, which we next define. If the motor distribution (Eq. 2) is tightly concentrated at the convergence zone (small *σ*), Eq. 4 is a good approximation for the behavior of the system for the points: 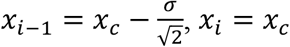, and 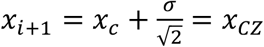

We can estimate values of the velocity for each of these points. The velocity at the convergence zone *x*_*CZ*_ is approximately zero because motor-based forces drive flow inward toward the convergence zone (*ν*_=+1_ = *ν*(*x*_*CZ*_) = 0). In turn, the velocity at the point 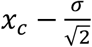 is estimated by noting that internal forces in a viscous fluid dissipate over a characteristic distance 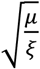 obtained by solving the equation: 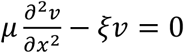. Based on this, velocity as a function of position is given by considering the effects of substrate adhesion, internal viscosity, and the spatial profile of active motors and is written as 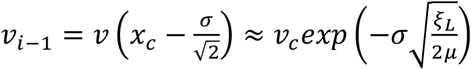.

To estimate the stresses associated with cellular deformation, we define *ν*_*i*_ = *ν*(*x*_*c*_) ≡ *ν*_*c*_ and put our estimates of *ν*_*i*−1_ and *ν*_*i*+1_ into Eq. 4. This yields the expression: 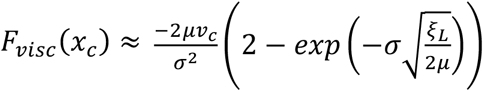, which in the limit 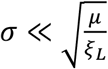 simplifies to:

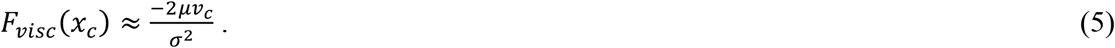

In plain language, Eq. 5 states that the local stress (e.g., tension) generated by cellular stretching or contraction rises as viscosity or deformation rate increases and decreases when the width of the motor distribution expands.

To determine the velocity of motion at the site of peak force generation, *ν*_*c*_, we determine the balance of stresses at *x*_*c*_, which is given by *F*_*m*_(*x*_*c*_) + *F*_*visc*_(*x*_*c*_) = *ξ*_*L*_*ν*_*C*_. By substitution, we rewrite this as 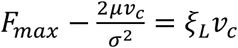 and solve for *ν*_*c*_:

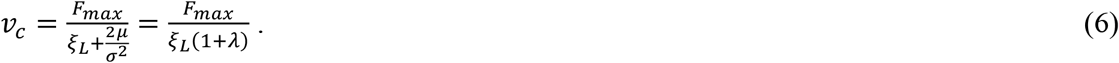

To abstract this equation in a way that allows consideration of the shape of the velocity profile based on the relative magnitudes of force generation, viscosity, and adhesion, we define a dimensionless parameter 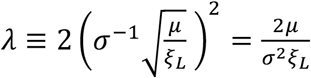. This equation indicates that *λ* measures the relative contributions of internal viscosity and external traction, defined by the ratio of the characteristic dissipation length, 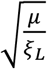, and the width of the motor distribution, *σ*.

Combining Eq. 6 with the expressions for viscous stress (Eq. 5) and traction stress yields:

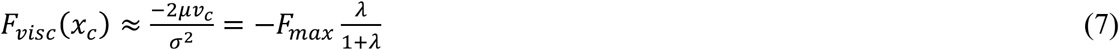

and

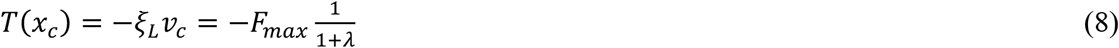

Importantly, equations 7 and 8 indicate that the relative magnitudes of viscous stress and traction stress at *x*_*c*_ are determined by a single parameter, *λ*. When *λ* is small, *F*_*visc*_(*x*_*c*_) approaches zero (Eq. 7) and *T*(*x*_*c*_) approaches –*F*_*max*_ (Eq. 8) meaning that motor-based forces are balanced and dissipated primarily by external traction (**Fig. 4C**). In this limit, cytoskeletal flow falls off rapidly with distance from the convergence zone (**Fig. 5A, B**). In contrast, for large *λ, F*_*visc*_(*x*_*c*_) approaches −*F*_*max*_ (Eq. 7) and *T*(*x*_*c*_) approaches zero (Eq. 8), meaning that motor-based forces are dissipated internally (**Fig. 4C**), giving rise to cytoskeletal flow with an approximately uniform velocity toward the convergence zone (**Fig. 5C**). Numerical solutions to Eq. 1 (data points in **Fig. 4C**) agree well with the analytical approximations (Eqs. 7 and 8) (solid lines in **Fig. 4C**).

**Figure 5.**
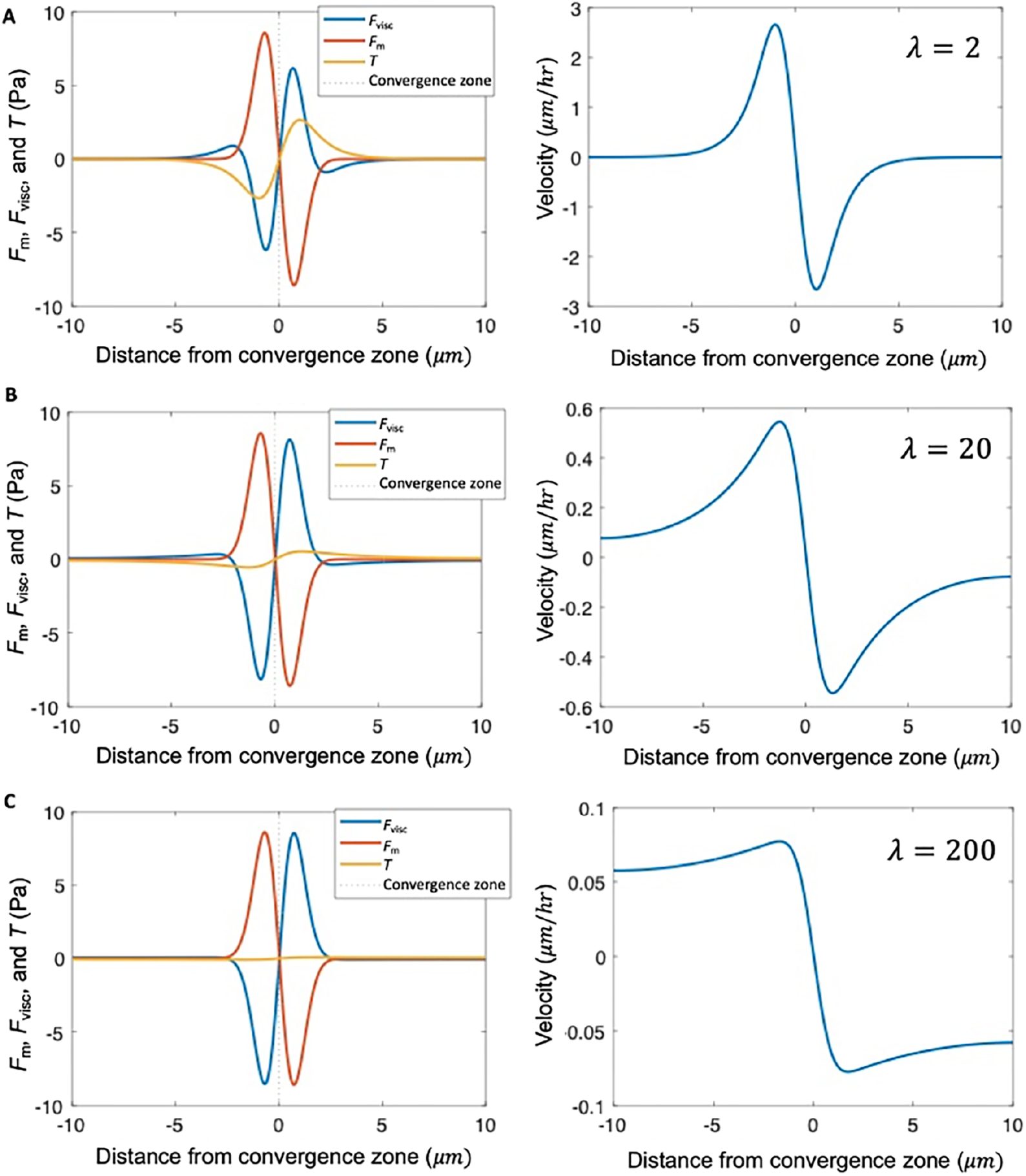
*λ* describes the shape of the velocity profile. Stress distribution and velocity profile calculated numerically from Eq. 1 for different levels of adhesion strength relative to viscosity, characterized by dimensionless parameter *λ*, for hypothetical cells with uniform viscosity *μ* and uniform adhesion strength *ξ* ≡ *ξ*_*L*_ = *ξ*_*R*_ **(A)** Balance of internal stress and traction (left) and flow field (right) for 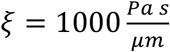, and 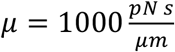, corresponding to characteristic length 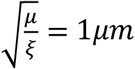 and *λ* = 2. **(B)** Same as (A), except with 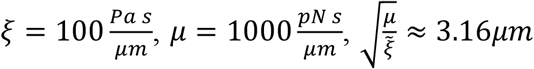 and *λ* = 20. **(C)** Same as (A), except with 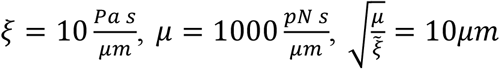, and *λ* = 200. Other input parameters for all panels: *L* = 20*μm, x*_*CZ*_ = 10*μm, σ* = 1*μm*, 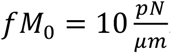.

### Approach for fitting experimental data

Putting these observations together, we can fit flow data from different cell types with numerical solutions to Eq. 1 to estimate the relative strength of adhesions on the left and right of the convergence zone, the maximum force generated across the convergence zone, and the ratio of viscosity over adhesion. Our strategy for constraining mechanical parameters of the model using experimental data is: (i) Set the location of the convergence zone, *x* = *x*_*CZ*_, to match the location where the magnitude of the experimental flow gradient is maximized; (ii) Set the motor distribution width, *σ*, to match the extrema of the second derivative in the experimental flow pattern; (iii) Tune the right-to-left adhesion ratio, 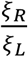, so that the numerical prediction matches the experimental measurement for the ratio of flow speed to the right and left of the convergence zone; (iv) Tune the ratio,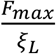, so that the rate of anterograde flow toward the convergence zone from the left agrees with experimental measurement; and (v) tune the ratio,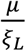, to match the measured velocity profile *ν*(*x*), left of the transition zone. Numerical solutions were obtained using finite difference numerical methods with custom code written in MATLAB. A summary of parameters for which the model approximately reproduces these key experimental observations is shown in **Table 2**.

**Table 2:** Parameters for model fits in Figs. 6 and 7.

| **Motility type** | **Motor distribution width** | **Adhesion strength ratio** | **Motor strength over left adhesion ratio** | **Viscosity over left adhesion ratio** | **Order of magnitude for dimensionless parameter** |
| --- | --- | --- | --- | --- | --- |
| | $\sigma \text{ } (\mu m)$ | $\frac{\xi_R}{\xi_L}$ | $\frac{F_{max}}{\xi_L} \left( \frac{\mu m}{s} \right)$ | $\frac{\mu}{\xi_L} (\mu m^2)$ | $\lambda \equiv \frac{2\mu}{\sigma^2 \xi}$ |
| Cytokinesis | 1.0 | 1.0 | 1.5 | $10^3$ | $\sim 10^3$ |
| Amoeboid migration | 9.0 | $4.0 \times 10^3$ | 1.52 | $5.4 \times 10^4$ | $\sim 10^3$ |
| Mesenchymal migration | 4.7 | 6.5 | 1.65 | $5.0 \times 10^2$ | $\sim 10^1$ |
| Neural migration | 1.0 | $6.0 \times 10^4$ | $3.2 \times 10^4$ | $4.5 \times 10^7$ | $\sim 10^8$ |
| Axon outgrowth | 1.0 | 1.2 | 1.0 | $2.9 \times 10^3$ | $\sim 10^4$ |

### Modeling specific modes of cell motility

A central prediction of our active fluid model is that cells that are symmetric in terms of their viscosity and adhesion distributions exhibit constant flow toward the convergence zone but are stationary. In contrast, asymmetries in viscosity or adhesions result in cell locomotion. To examine the flow and motility of specific types of cells, we took 2D timelapse images (Schaub et al., 2007) and created 1D kymographs along the flow axis to measure the velocity profile. Starting with cytokinesis, we set the model such that the convergence zone is centered and adhesion strength is small and identical on both sides. This generates a stationary symmetric inward flow pattern that tightly fits the experimental data (**Fig. 6A**). In contrast, by introducing a large magnitude of substrate adhesion under the leading edge while keeping substrate adhesion under the cell body very low, the model reproduces slow leading-edge retrograde flow in conjunction with rapid anterograde flow in the cell body characteristic of rapid amoeboid migration (**Fig. 6B**). In turn, by lowering the front/rear adhesion ratio and shifting the convergence zone location, the model produces flow patterns and motility that fit mesenchymal migration (**Fig. 6C**). In both cases, the positive velocities generated by the model across the left-hand side match the forward motion of the cell. A side-by-side comparison of the experimental data indicates that the rates and overall flow patterns that occur during cytokinesis, amoeboid, and mesenchymal motility are nearly identical. The modeling suggests they differ primarily in the position of the convergence zone and the relative strength of adhesions on the left and right-hand sides.

**Figure 6.**
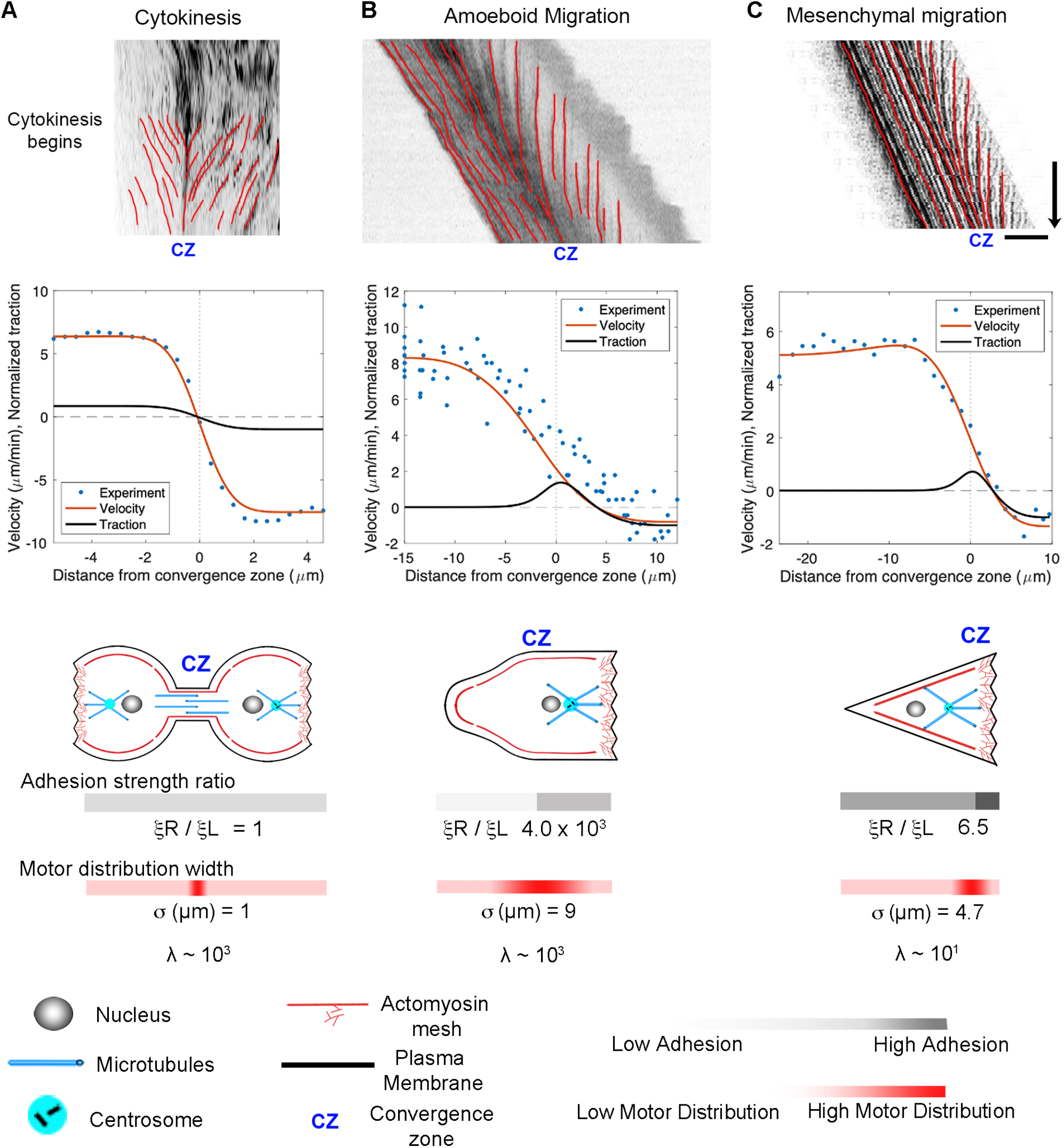
Adhesion asymmetries and convergence zone location explain differences between cytokinesis, amoeboid migration, and mesenchymal migration. (A) Kymographs illustrating flow during cytokinesis, (B) amoeboid migration, and (C) mesenchymal migration. In the graphs, the blue dots represent experimental data, and the red and black lines show numerical predictions of the flow and corresponding traction distribution. The model fits for adhesion strength ratio, motor distribution, and *λ* are shown below; bar = 10 *μ*m, arrow = 1 min.

### Modeling of neuronal migration and axonal outgrowth

The growth cone, alliteratively, has been called “a leukocyte on a leash” (Bray and White, 1988; Pfenninger, 1986), and several groups have noted a likeness between migrating cells and advancing growth cones (Aberle, 2019; Miller and Suter, 2018; von Philipsborn and Bastmeyer, 2007). It is well-accepted that migrating neurons exhibit anterograde flow in their leading process, consistent with components moving synchronously in the direction of motility (Guan et al., 2007; He et al., 2010; Hutchins and Wray, 2014; Minegishi et al., 2018). At the same time, the P-domain of the growth cone maintains its width through a balance of actin assembly at the leading edge and disassembly at the convergence zone. Thus, neuronal migration resembles mesenchymal motility, differing primarily in cell shape rather than the underlying sub-cellular motion. Likewise, a series of studies have demonstrated that *Drosophila, Aplysia, Xenopus*, Rat, and Chicken peripheral neurons lengthen through the bulk forward flow of materials in the distal axon (Athamneh et al., 2017; Lamoureux et al., 2010b; Reinsch et al., 1991; Roossien et al., 2013). This suggests that axonal outgrowth and neuronal migration are highly similar, differing primarily in the motion of the proximal region of the axon attached to the cell body.

Motivated by these observations, we extend the model to describe the flow patterns and internal force distributions in migrating neurons and elongating axons (**Fig. 7**). To simplify, we assume that forces induced by motor proteins along the axon are balanced and that contractile force generation in the growth cone dominates. To model this, the axon is treated as a 1D domain extending from its proximal edge (*x* = 0) to the growth cone tip (*x* = *L*) with motor proteins centered at the growth cone transition zone, *x* = *x*_*CZ*_ as shown in the schematic (**Fig. 7A**). The motors simultaneously drive retrograde flow in the growth cone and pulls material forward in the leading process and distal axon. To model neuronal migration, we created kymographs from time-lapse movies showing the motion of alpha-actinin-GFP (He et al., 2010) to track the velocity of actin filaments along the length of the leading process (top panel) and over the P-domain of the growth cone (bottom panel). Data from these examples were used for the analysis in the graph noted as Experiments 1 and 2 (**Fig. 7C**). The velocity profile indicates that retrograde flow occurs at a modest velocity across the P-domain of the growth cone, and the leading process and cell body advance at a uniform rate over distance. Fitting the model to the data (**Fig. 7D**) suggests growth cone adhesion is substantially higher than leading process adhesion,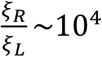, which is consistent with reports of the formation of strong adhesions in front of the cell body that generate traction that pulls the components of migrating neurons forward, coupled with the removal of adhesive elements towards the rear (Solecki, 2012). Tuning the model parameters to fit the data results in a narrow convergence zone (i.e., small *σ*) and a high viscosity relative to adhesion. These conditions indicate that 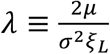 is large (∼10^8^), consistent with the experimental data showing a roughly constant velocity of motion across the cell body and along the length of leading process of migrating neurons (**Fig. 7C**).

**Figure 7.**
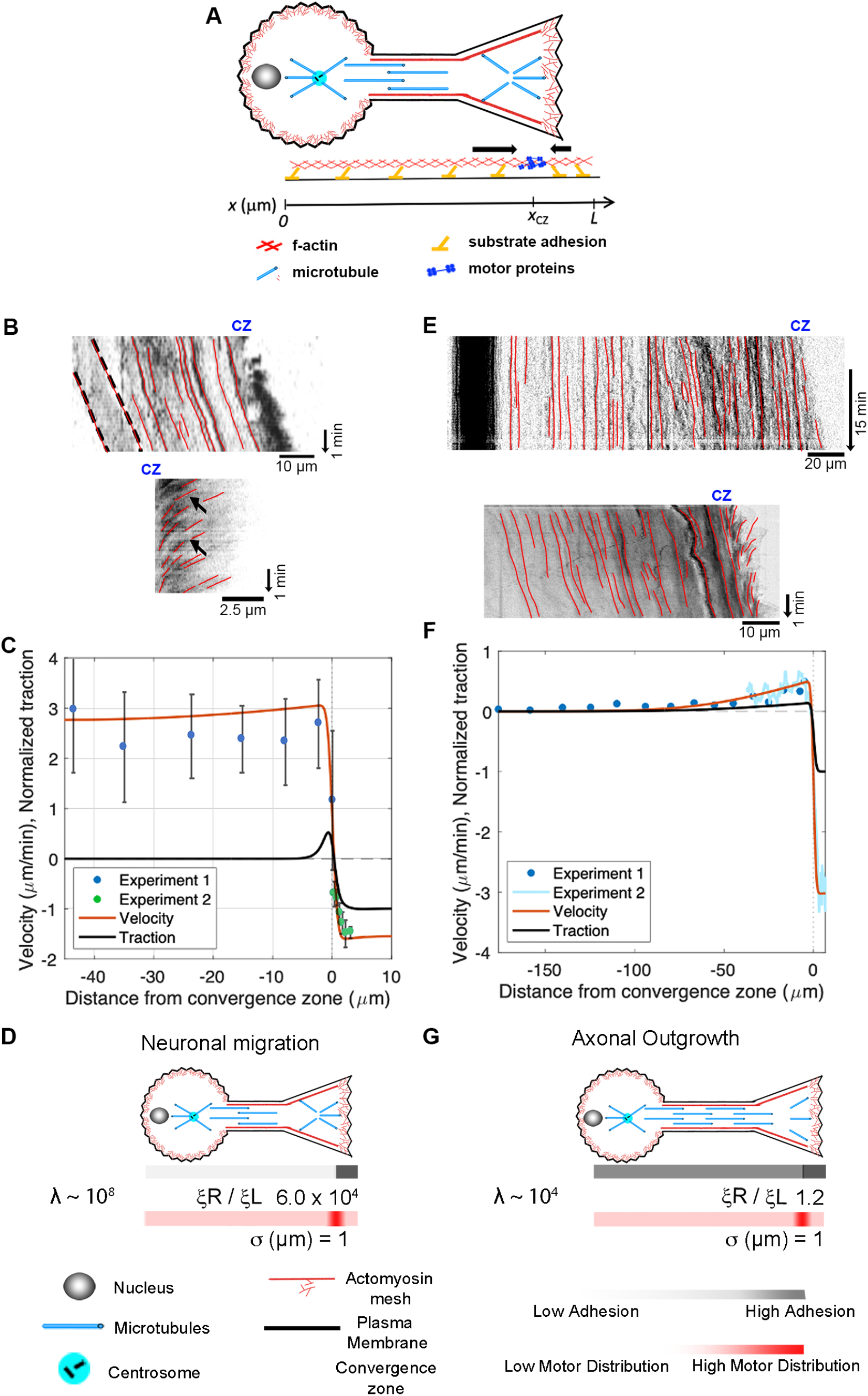
Differences between neuronal migration and axonal outgrowth are explained by leading process / axonal adhesion strength. (A) Schematic of growth cone motility. (B) Kymographs of a migrating neuron using alpha-actinin-GFP to track actin filaments along the leading process (top) and across the growth cone (bottom). (C) Numerical predictions of the flow pattern (red line) and corresponding traction distribution (black line) with experimental data shown as blue and green dots. (D) Numerical fits for neuronal migration indicate that the ratio of adhesion between the growth cone and the leading process and *λ* are both large. (E) Kymographs of sensory neuron outgrowth using docked mitochondria to track flow along the length of a axon (top) and phase images to assess retrograde flow across the growth cone (bottom). (F) Data from these experiments, with predictions of the velocity profile (red line) and traction forces (black line). (G) Model agreement with the experimental flow pattern is achieved with λ ∼ 10^4^ and a front-to-back adhesion ratio on the order of 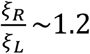.

Next, we apply our mechanistic framework to axonal outgrowth. Growing axons exhibit bulk flow distally that matches the rate of growth cone advance, but materials are stationary in the proximal axon. The top kymograph (**Fig. 7E**) shows the flow of docked mitochondria (top) in a sensory neuron undergoing axon outgrowth (Athamneh et al., 2017). The dark band on the left indicates that the cell body is stationary. The red lines along the axon illustrate that material in the proximal axon is also stationary relative to the substrate, while forward advance occurs in the distal axon. The bottom kymograph, created from phase images, shows forward flow in the distal axon and retrograde flow across the growth cone. Data from these experiments are shown as Experiments 1 and 2 in the graph below (**Fig. 7F**). Model agreement with the experimental flow pattern is achieved for a front-to-back adhesion ratio on the order of 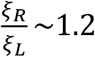 (**Fig. 7G**). This suggests that neurons switch from migration (**Fig. 7D**) to elongation (**Fig. 7G**) by increasing adhesion under the axon and cell body (Calof and Lander, 1991; Minegishi et al., 2018; Solecki, 2012).

## DISCUSSION

The primary advance presented in this study is the development and application of an active fluid model that describes the flow of cytoskeletal elements during various cellular processes, including cytokinesis, amoeboid migration, mesenchymal migration, neuronal migration, and axonal outgrowth (**Fig. 6, 7**). The key finding is that differences in motility and flow patterns can be explained by variations in adhesion strength on the leading and lagging sides of cells, viscosity, and the positioning of primary sites for contractile force generation. To the best of our knowledge, this study is the first to create a general model that relates cytokinesis, various forms of cellular motility, and axonal outgrowth. It suggests that the biophysical mechanism of axon outgrowth is closely related to mechanisms that drive cell crawling and cytokinesis.

### Working towards a global active fluid model of neuronal mechanics

Interest in neuronal mechanics is flourishing (Franze, 2020; Ghose and Pullarkat, 2023; Miller and Suter, 2018; Oliveri and Goriely, 2022; Raffa, 2023). In this context, active fluid models provide a useful framework for describing the flow of materials where force-generating mechanisms are embedded throughout a material (Julicher et al., 2007; Mogilner and Manhart, 2018). These models treat materials as viscoelastic fluids that contain motors producing extensile and contractile forces (de Rooij et al., 2017). Doing so updates classic models by providing a new solution to how cells simultaneously maintain a continuous flow of materials and constant tension. To briefly review, when the biophysical properties of neurons were initially modeled (Dennerll et al., 1989), axons were viewed as passive viscoelastic solids: in essence, a single spring connecting the cell body and growth cone. This fits with the ideas that the growth cone pulled the axon forward, generating a rest tension (Lamoureux et al., 1989) and earlier studies suggesting microtubules do not move out of the cell body by bulk flow (Bamburg et al., 1986; Miller and Joshi, 1996; Okabe and Hirokawa, 1990). Coupled with an understanding that growth cones pull (Lamoureux et al., 1989), the appeal of this model was that it explained how axons maintain a constant rest tension without materials flowing toward the growth cone. Later, when bulk flow of microtubules was observed in the distal axons of frog neurons (*Xenopus laevis*) (Reinsch et al., 1991) and in other species (Athamneh et al., 2017; Miller and Sheetz, 2006), treating the axon as a viscous-elastic solid became problematic because solids do not flow. The first step in addressing this problem was to treat axons as a passive viscoelastic fluid bound to the substrate through adhesions instead of a solid to explain the forward flow of material toward an actively pulling growth cone (O’Toole et al., 2008). Yet, because tension dissipates over distance when passive fluids interact with a substrate, a shortcoming of this model was that it did not explain how axonal tension is maintained far from the growth cone. This issue was solved by the addition of embedded motors throughout an axon, modeled first as a viscoelastic solid and then later as a Maxwell fluid (Bernal et al., 2007; de Rooij et al., 2018; Recho et al., 2016). The second issue was that early models treated the growth cone as a single force vector (O’Toole et al., 2008). This was addressed by combining measurements of the viscoelastic properties of growth cones with flow maps of actin motion across the growth cone to infer patterns of subcellular force generation (Betz et al., 2011). Later, experimental verification that tension is generated in axons by actomyosin and that these contractile forces are countered by dynein-driven microtubule sliding provided a molecular foundation for the active nature of axons (O’Toole et al., 2015; Roossien et al., 2014; Tofangchi et al., 2016). Incorporating these ideas into an active viscoelastic fluid model helps to explain why axons act as solids over short time scales (Bernal et al., 2007) but fluids over long timescales (de Rooij et al., 2017); and provides a foundation for modeling an extensile core of microtubules (Roossien et al., 2014) around a contractile shell of actomyosin (de Rooij et al., 2018; Recho et al., 2016). The contribution of the work presented here is the development of a single model of both the axon and the growth cone that fits experimental data describing the flow of material from the tip of the P-domain to the neuronal cell body (Athamneh et al., 2017). Thus, we suggest this is the first global, though highly simplified, active fluid model of axonal outgrowth. Given the history of this problem, the surprising aspect, at first, is that by modifying parameters in a model initially developed for axonal outgrowth (**Figs. 6 and 7**), the flow maps of cytokinesis, amoeboid migration, mesenchymal migration, and neuronal migration are well-fitted.

### Comparing Cytokinesis, Amoeboid, and Mesenchymal Motility

The overall morphology and cytoskeletal flow maps of cytokinesis and amoeboid migration share similarities that have long been appreciated (Bray and White, 1988; DeBiasio et al., 1996; Swann and Mitchison, 1958). In both, cells have a cylindrical morphology with a centralized convergence zone (**Fig. 2**) where the actomyosin-based cell cortex flows at approximately 5 *μ*m/min (**Fig. 6 A, B**). The modeling suggests a major difference between flow during cytokinesis and amoeboid migration is that the convergence zone is confined to a narrow region during cytokinesis but is roughly 10-fold wider during amoeboid migration (**Table 2**). At a biological level, this is consistent with Rho-mediated actomyosin contraction being activated in a precise zone at the cell equator via MgcRacGAP during cytokinesis (Miller and Bement, 2009), but having widespread activation, except at the leading edge where Cdc42 inhibits Rho, across amoeboid cells, (Yang et al., 2016). The second difference is that flow is relatively symmetric during cytokinesis but is highly asymmetric during amoeboid migration (**Fig. 6**). Fitting this data suggests adhesions are balanced during cytokinesis but differ by 1000-fold during amoeboid migration (**Table 2**). A plausible interpretation of this data for ameboid cell migration, as noted previously (Liu et al., 2015) and seen in **Fig. 2B**, is that the cell body pushes into the sides of the channel, but the uropod is narrower and acts as a weakly adhering passive dragged body.

While cells undergoing amoeboid and mesenchymal migration have dramatically different morphologies, the underlying cytoskeletal flow maps are similar (**Fig. 6B, C**). Based on our modeling, when transitioning from amoeboid to mesenchymal migration, overall levels of adhesion increase by a factor of 100. At the same time, the ratio of adhesion between the front and back of the cell decreases by a factor of 1000 (**Fig. 6 and Table 2**). This aligns with observations that mesenchymal cells possess a weak gradient of strong substrate adhesions, while amoeboid migration involves a strong gradient of weak non-specific substrate interactions (Bergert et al., 2015; Paluch et al., 2016; Parsons et al., 2010). These differences arise because, during mesenchymal migration, relatively strong substrate adhesions are found across the cell, while in amoeboid migration, the leading edge pushes strongly against the substrate in confined channels, but the trailing uropod interacts very weakly. In this context, the model predicts that the reason traction forces generated during amoeboid migration are particularly low is that the back of the cell does not oppose the front.

### Mesenchymal and Neuronal Migration

While migrating neurons and non-neuronal cells undergoing amoeboid or mesenchymal migration have dramatically different morphologies, the overall flow profile appears similar (**Figs. 6B, C, and 7B, C**). In all three cases, there is a convergence zone towards the front of the cell, a region of retrograde actin flow at the leading edge and forward advance of the cell body. Nonetheless, there are rather dramatic differences in the biophysical parameters (**Table 2**). In particular, the length of the contractile zone in neuronal migration is roughly 5-fold shorter than in non-neurons, reflecting the fact that growth cones of migrating cerebellar neurons span roughly 5 *μ*m, while the leading edge of fish keratocytes is roughly 20 *μ*m (**Fig. 2C, D**). Furthermore, *λ* (**Table 2**), which controls how rapidly a cell stretches over distance as it is being pulled across a substrate, is eight orders of magnitude higher in migrating neurons than in mesenchymal cells. Here, an important underlying factor may be that migrating neurons decrease adhesion strength (*ξ*) under the axon to decrease frictional forces. As adhesion regulation is recognized as being critical for neuronal migration, and the signaling pathways and cell adhesion molecules are relatively well understood (Calof and Lander, 1991; Minegishi and Inagaki, 2020; Solecki, 2012), an experimental comparison of adhesion strength between cells undergoing mesenchymal and neuronal migration would both help to test the predictions of this model and to understand the similarities and differences between different modes of cell crawling.

### Switching From Neuronal Migration to Axon outgrowth

Migrating neurons and neurons undergoing axonal outgrowth have very similar morphologies, but the mechanisms underlying neuronal migration and axon outgrowth have generally been thought to be unrelated. By comparing the flow patterns in the distal axon, we show here that the mechanics of growth cone advance are similar. In both cases, under the P-domain of the growth cone retrograde flow pulls the substrate rearwards, while material along the axon flows forward at the rate of growth cone advance. Where they differ is that flow velocity along the length of migrating neurons is relatively constant, but during axonal outgrowth, it drops to zero towards the cell body (**Fig. 7C, D**). The model predicts this decline in bulk flow occurs because the ratio of viscosity over adhesion 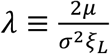 along the leading process is four orders of magnitude higher during neuronal migration than during axonal outgrowth (**Table 2**). Noting endocytosis-mediated de-adhesion under the cell rear (Shieh et al., 2011; Solecki, 2012) coupled with the formation of strong adhesions under the growth cone (Minegishi et al., 2018) is required for neuronal migration, the model is consistent with experimental data indicating that the transition from neuronal migration to axonal outgrowth occurs as the result of changes in adhesion under the cell body and leading process. Collectively, this suggests that the primary difference between neuronal migration and axonal outgrowth lies in the behavior of adhesions under the leading process and cell body, rather than the mechanics of the growth cone.

### Limitations

The primary limitation in developing a general biophysical model is that complex details, such as the interplay between extensile and contractile forces, are reduced to a simplified set of generalized parameters. In these terms, the model, as written, has three primary limitations. The first is that it does not explicitly model the balance of forces associated with actin filament assembly against the plasma membrane (Craig et al., 2012), extensile forces associated with MT sliding (Lu et al., 2013; Roossien et al., 2014), observations of multiple contractile zones in cells (Jiang et al., 2015), nor how the system responds elastically to abrupt changes in force (Bernal et al., 2007). The first two issues can be addressed by adding additional convergence or extensile force-generating zones in the computational model and including a term for membrane tension. The second can be addressed by adopting an active Maxwell fluid approach (de Rooij et al., 2017) by expanding the viscosity term (*η*) into elastic (*E*) and time scale (*τ*) components (*η* = *Eτ*) and rewriting the equations to be time-dependent (Betz et al., 2011). Nonetheless, for the purposes of this work, because we are focused on steady-state conditions, where there is only one apparent contractile zone in the experimental data for each of the modes of motility we consider, the approach we use seems to strike a good balance between being complex enough to model the experimental data while maintaining a generality that allows comparisons between different cell types.

The second major challenge in developing models that unite modes of cell motility is a lack of systematic measurements across diverse cell types that generate output at the subcellular level in meaningful physical units (e.g., *N, Pa-sec, Pa*, or *N/m*) of local force generation, viscosity, elasticity, and adhesion. While the use of adhesion ratio and the *λ* term allows non-dimensional modeling of cell motility, the direct way to test model predictions will be to develop collaborations to measure these parameters across the modes of motility. Recent progress in this field has been exciting, and the combination of clever experimental approaches (Ghose and Pullarkat, 2023), sophisticated biophysical models (Oliveri and Goriely, 2022), and powerful molecular genetic tools (Yasunaga et al., 2019) has the promise of moving cellular biophysics toward a unified theory of eukaryotic cytokinesis, cell crawling, and axon outgrowth.

## CONCLUSION

It’s long been appreciated cell crawling is a continuation of cytokinesis (Bray and White, 1988; DeBiasio et al., 1996; Swann and Mitchison, 1958), mesenchymal migration and amoeboid migration are points on a continuum (Liu et al., 2015), and that growth cones resemble crawling cells (Bray and White, 1988; Pfenninger, 1986). The model presented here provides a simple means to understand how they are related. In each case, there is a primary contractile zone that drives flow, and differences in subcellular adhesion patterns dictate intracellular flow and motility rates. Noting that the signaling pathways and effectors that control these activities (e.g., Rho, Rac, NMII, and integrins) are well conserved raises the possibility that the reason why cytokinesis, cell crawling, and axon outgrowth can be described by modifying parameters in a simple model is that they are evolutionarily related.

## METHODS

### Acquisition of Experimental Data and Permissions

Flow rates for cytokinesis were obtained by importing an image of Fig. 1B from (Singh et al., 2019) and extracting the data points for flow speed as a function of % cell length using ImageJ. The still image and kymograph for **Fig. 2A** in this manuscript were obtained by downloading Movie 1 from (Singh et al., 2019) and creating a kymograph in ImageJ. With permission from the Journal of Cell Science, Creative Commons Attribution License 4.

The figure panels for **Fig. 2B** and the flow rates for amoeboid migration were obtained by downloading the movie titled “Myosin dynamics at the cortex of a Walker cell migrating in a BSA-coated channel” from (Bergert et al., 2015). ImageJ was then used to create a kymograph and to measure subcellular flow. With permission from Springer Nature.

The figure panel and flow rates for mesenchymal migration for **Fig. 2C** were obtained by downloading “video01” from (Schaub et al., 2007), and then using ImageJ to create a kymograph and to measure subcellular flow. Because the time-lapse movie was short, 30 sec, we stacked and realigned the same kymograph six times to make the figure panel with the intent of making it easier to compare modes of motility visually. With permission from the author, Alexander Verkhovski, and from Mol. Cell Biol. Creative Commons Noncommercial Share Alike 3.0 Unported license.

The DIC figure panel in **Fig. 2D** was created by reformatting a portion of Fig. 6A in (He et al., 2010). The fluorescent image in **Fig. 2D** and the kymograph for neuronal migration were obtained by downloading “Movie_S3” and processing it in ImageJ. The estimate of retrograde flow velocity in the growth cone of migrating neurons shown in **Fig. 6B** was based on an analysis of the kymograph in Sup. Fig. 5B in (He et al., 2010). With permission from the author, Xiaobing Yuan, and the Journal of Neuroscience, Creative Commons Attribution Noncommercial Share Alike 3.0 Unported License.

Flow rates for axon outgrowth in the distal axon and growth cone in **Fig. 2E, F** and **Fig. 7D, E** were obtained from the source data used to generate Figs. 3 and 4 in (Athamneh et al., 2017). With permission from the author, Kyle Miller, and Science Reports Creative Commons Attribution License 4; J. Cell Science, CC-BY license.

## AUTHOR CONTRIBUTIONS

KM analyzed the experimental data, EC conducted the biophysical modeling, FO and SA constructed figures, and KM and EM wrote the manuscript and developed the figures.

## ACKNOWLEDGMENTS

Work related to this article was funded by NSF Research at Undergraduate Institutes Award 1915477 to Erin M. Craig and NIH (5R01MH094607-05) and NSF (Award number 1453799) to Kyle E. Miller. We are grateful to Steven Heidemann, Anthony Cognato, and Kahmina Ford for helpful discussions.

